# Target Enrichment Enhances the Sensitivity of Sanger Sequencing for BRAF V600 Mutation Detection

**DOI:** 10.1101/2021.08.14.456349

**Authors:** Qiang Gan, Andrew Fu, Fang Liu, Shuo Shen, Maidar Jamba, Wei Liu, Mike Powell, Aiguo Zhang, Michael Sha

## Abstract

BRAF is a serine/threonine protein kinase whose mutations lead to unregulated cell growth and cause different types of cancers. Since V600E is a major BRAF mutation and V600E detection as a companion diagnostic test (CDx) is stipulated in the labeling of the BRAF V600 inhibitors. Traditional Sanger sequencing cannot accurately detect mutations lower than 15% variant allele frequency (VAF) due to its limited sensitivity. Here we applied our patented XNA molecular clamping technology to modify Sanger sequencing template preparation by enriching the mutation population. We found that the use of our mutation-enriched template enhanced the analytical sensitivity of Sanger sequencing to 0.04% VAF. The method is verified to detect V600E, V600K, and V600R mutants and is validated for the known BRAF mutation status in clinical samples. Our streamlined protocol can be used for easy validation of the highly sensitive target-enrichment method for detecting BRAF V600 mutations using Sanger sequencing in clinical labs. In addition to BRAF V600 mutations, this method can be extended to the detection of other clinically important actionable mutations for cancer diagnostics.

## Introduction

BRAF is a serine/threonine protein kinase in the Ras/Raf/MEK/ERK pathway that regulates apoptosis and cell cycle progression (1,2). Mutations in RAF proteins lead to unregulated cell growth and cause different types of cancers due to the activation of several downstream pathways. Major RAF point mutations are in BRAF with rare occurrences in related ARAF and CRAF genes (3). The major BRAF-activating mutation V600E is a single nucleotide c.1799T>A substitution at exon 15, which leads to BRAF overexpression and a constitutive elevated activity (500-fold more activity compared to normal BRAF activity) (4). Another important BRAF point mutation, BRAF V600K with two nucleotide substitutions (c.1798_1799GT>AA), has a poorer prognosis compared to BRAF V600E (5). Other important BRAF V600 mutations that occur much less frequently include V600R, V600D, V600M and V600G (6).

In melanomas, 50% of cases carry BRAF mutations, out of which >90% of them are BRAF V600E mutations (7). BRAF V600K accounts for 5 to 6% of total mutations and BRAF V600D or V600R mutations are much less frequent. In lung cancer, BRAF mutations are found in 2–4% of lung adenocarcinomas, especially non-small cell lung cancer (NSCLC) (8). In colorectal cancer cases, 10% of patients harbor BRAF mutations, of which only the BRAF V600E mutations are associated with poor prognosis (9).

Different BRAF inhibitors have been developed and approved by the U.S. FDA for targeted therapy toward BRAF V600E or both BRAF V600E and V600K mutations. These inhibitors include vemurafenib, dabrafenib, encorafenib, and MEK inhibitors, such as binimetinib and cobimetinib (10, 11, 12). To initiate target therapy using BRAF V600 inhibitors, the FDA requires companion diagnostic tests (CDx) to be performed to confirm the mutation. Multiple *in vitro* molecular diagnostic tests, including Sanger sequencing, qPCR, allele-specific PCR (13), amplification refractory mutation system (ARMS) (14), COLD-PCR (15), HRM (16), pyrosequencing (17), mass spectrometry (18), XNA (Xenonucleic acid) molecular clamps (19, 20), droplet digital PCR (ddPCR) (21), and more recently NGS (22, 23, 24), are available for mutation testing, each with different levels of sensitivity.

Sanger sequencing is routinely used to compare and evaluate newer analytical methods, although it falls short in terms of use with CDx assays because of its inability to provide the low limit of detection (LoD) required for them. To improve the analytical sensitivity of Sanger sequencing for the BRAF V600 mutation assay, we leveraged the XNA molecular clamps developed at DiaCarta to enrich the mutant population in a normal wildtype background. XNA is a synthetic DNA analog that can selectively block amplification of the wildtype sequence but allows a low-frequency mutation sequence to amplify in a PCR reaction. The mutation enrichment significantly improves assay sensitivity. We have shown that this high-sensitive, target-enrichment method for Sanger sequencing is simple and can be easily validated for single BRAF V600 mutation in clinical labs for companion diagnostic testing.

## Materials and Methods

### Reference materials and clinical samples

The reference standard for wildtype human genomic DNA, BRAF V600E, V600K, and V600R mutations were purchased from Horizon Discovery. The mutant DNA was mixed with the wildtype DNA at different VAF for limit of detection determination. The clinical FFPE samples were purchased from BioChain (Hayward CA).

### DNA preparation and quantification

The genomic DNA from clinical FFPE samples was extracted using QIAamp DNA FFPE Tissue Kit from Qiagen and quantified using Qubit from Thermo Fisher Scientific.

### BRAF V600 XNA synthesis

BRAF V600 XNA was designed, synthesized, and purified at DiaCarta Inc (Richmond, CA). The synthesized XNA was characterized by HPLC and mass spectrometry.

### BRAF V600 mutation enrichment and detection by XNA-based qPCR

For the qPCR test, different concentrations of BRAF V600 XNAs (8 μM to 0.25 μM for optimal XNA concentration) or a different mutant over wildtype population ratio (VAF) (5% to 0% for LoD assay) were used. Briefly, the 10 μl real-time PCR reaction contains 5 μl 2X Master Mix, 2 μl primer and probe mix, 1 μl BRAF V600 XNA, 2 μl DNA sample (10 ng) or control. The RT-PCR reaction was performed using the Bio-Rad CFX384 instrument under the following conditions: 98°C for 5 minutes followed by 50 cycles of 95°C for 20 seconds, 70°C for 40 seconds, 64°C for 30 seconds, and 72°C for 30 seconds before storage at 10°C.

### BRAF V600 mutation enrichment and detection by Sanger sequencing

For BRAF V600 mutation enrichment for Sanger sequencing, the enrichment PCR was performed as follows: the enrichment 20 μl-PCR reaction contained 1μl (10 ng) template DNA, 2 μl primers mix, 2 μl BRAF V600 XNA, 10 μl 2X Master Mix, and 5 μl nuclease-free water. The reaction was performed under the following conditions: 98°C for 30 seconds followed by 37 cycles of 98°C for 15 seconds, 60°C for 1 minute and 72°C for 15 seconds. The tubes were incubated at 72°C for another 2 minutes before storing at 10°C. The PCR product was purified with 1.5x ratio KAPA pure bead (Roche Cat# KK8002) and then eluted into water. After BRAF V600 mutation enrichment by XNA-based standard PCR, the samples were sent to Sequetech for Sanger sequencing.

### Validation of the BRAF V600 mutation-enrichment coupled Sanger sequencing

For validation of the BRAF V600 mutation-enrichment coupled Sanger sequencing, three kits from AdvancedSeq, including Exo-Alp PCR Cleanup Mix, SupreDye™ Cycle Sequencing Kit, and SupreDye™ XT Purification Kit, were used after enriching the BRAF V600E mutation as described in the user manuals. Briefly, to a 10 μl BRAF mutation enrichment PCR reaction, 4 μl Exo-Alp PCR Cleanup Mix was added and incubated at 37°C for 15 min and 80°C for 15 min. The samples were diluted 8-fold and the sequencing reaction was set up in a microtiter plate as follows: 0.4 μl SupreDye v3.1 Cycle Sequencing Ready Mix, 1.8 μl 5X Sequencing Reaction Buffer, 2.0 μl template (diluted PCR product sample, or positive, or negative, or no template control), 1.0 μl BRAF forward primer or reverse primer (5 μM), and 4.8 μl nuclease-free water. The cycle sequencing reaction was set up on the PCR instrument (AB 9700 GeneAmp) with the following conditions: 95 °C for 30s followed by 95 °C for 10s, 50 °C for 5s, 60 °C for 2min for 30 cycles. The sequencing reaction was purified by SupreDye XT Purification Kit as follows: 10 μL SupreDye XT resin and 45 μL SupreDye XT solution were added to the 10 μL sequencing reaction above. The plate was sealed and vortexed for 30 minutes at room temperature before loading the samples to AB 3730XL Genetic Analyzer for Sanger sequencing.

## Results

### BRAF V600 mutation can be enriched and detected by qPCR

First, we used qPCR to see if the BRAF V600E mutation is enriched and can be detected by qPCR. Our results showed that the BRAF mutation was selected to be amplified at various allele frequencies while the wildtype sequence amplification was blocked (Figure 1). The higher the VAF, the earlier was the amplification cycle when the BRAF V600E mutation could be detected. At 1μM XNA concentration, the BRAF V600E mutation could be detected at as low as 0.0625% VAF with a 100% correct call, corresponding to 1-2 copies of mutant DNA in 10 ng total gDNA input per test in the PCR template or reaction.

**Figure 1.**
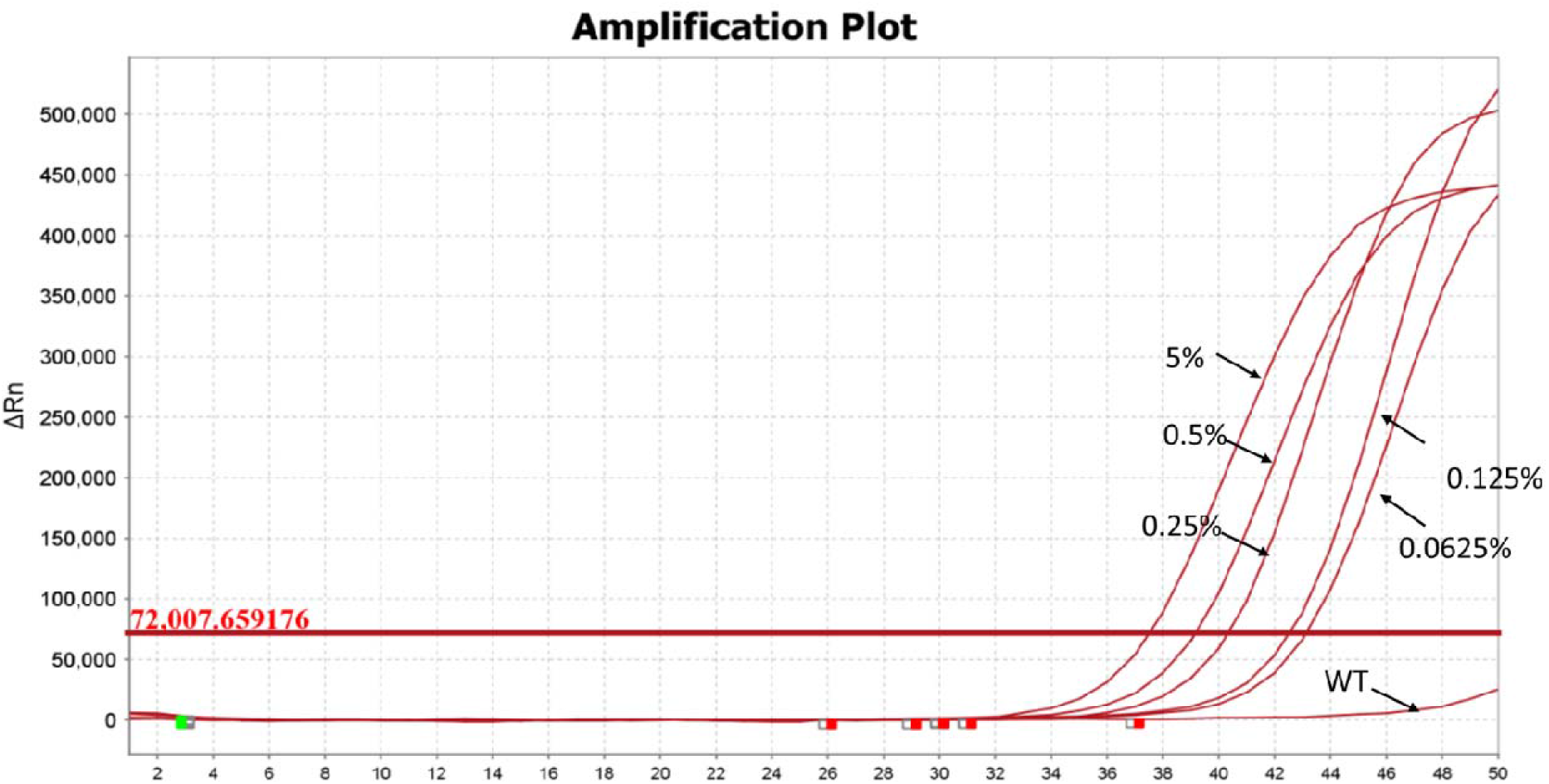
The qPCR amplification plot for BRAF V600E mutation detection in the presence of 2 μM XNA using templates with different VAF from 0% (WT) to 5%.

### BRAF V600 enrichment to improve the analytical sensitivity of Sanger sequencing

We prepared the BRAF V600E template by mutation-enriching PCR and used Sanger sequencing for mutation detection. The test workflow is shown in Figure 2a and the locations of the XNA and primers used for PCR and sequencing are depicted in Figure 2b.

**Figure 2.**
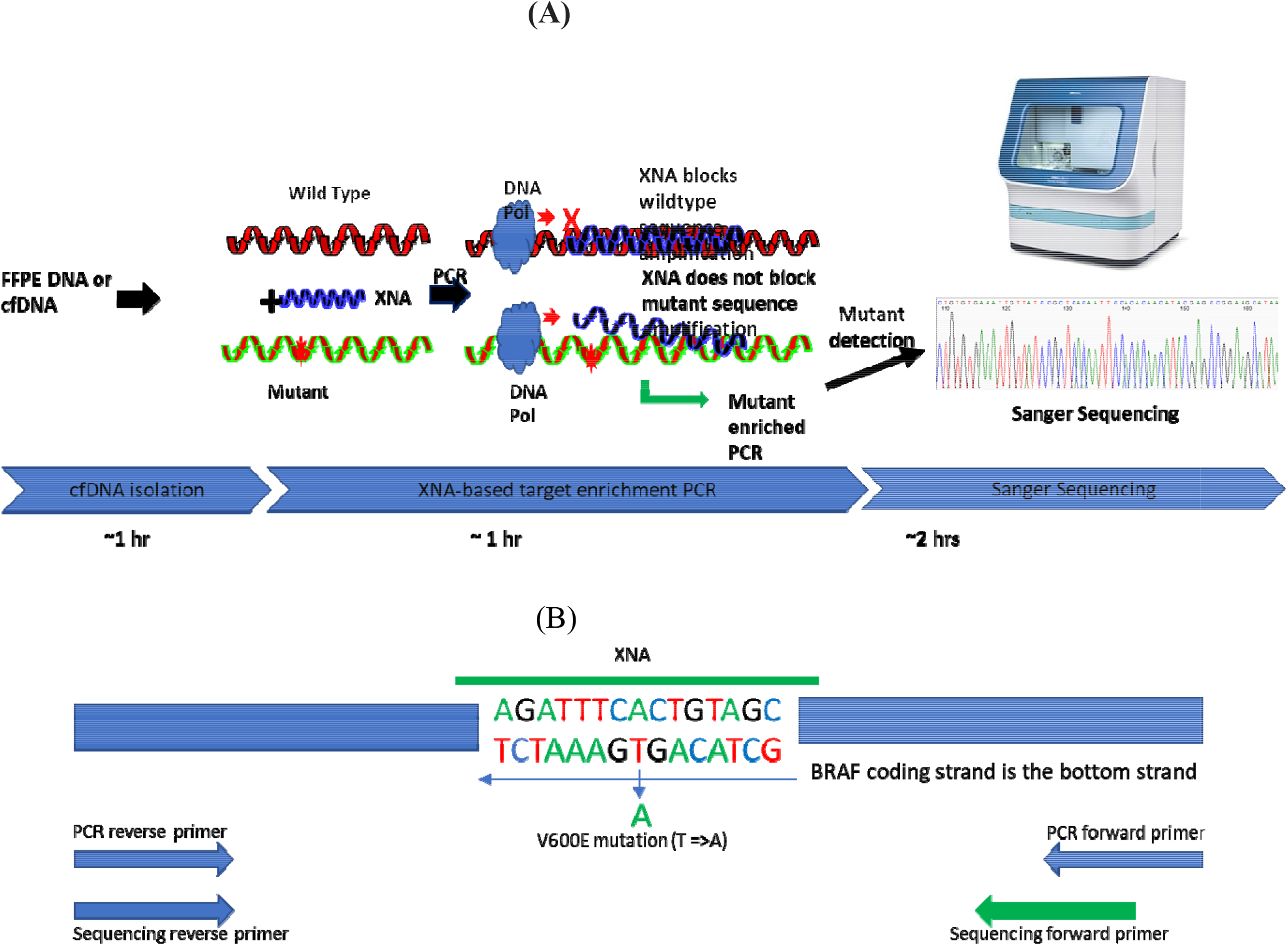
(A). The workflow for target-enrichment using XNA technology and subsequent Sanger sequencing. The XNA molecular clamps block the wildtype sequence amplification and selectively amplifies the mutant sequence, thereby increasing the sensitivity of the Sanger sequencing. (B). The positions of primers and XNA flanking or at the BRAF V600E mutation site are shown. These primers and XNA are used in the enrichment PCR reactions and subsequent cycle sequencing reactions.

To test the clamping effect of XNA concentration on mutation detection we performed experiments with different XNA final concentrations at 0.625% VAF. We found that when BRAF V600 XNA reached the final concentration at 500 nM only the mutation sequence peak, but not the wildtype sequence peak, was detected with Sanger sequencing (Fig. 3a). These data indicate that 500 nM final concentration is an appropriate concentration to detect low VAF of BRAF V600 mutations. To determine the limit of detection (LoD), we performed experiments with a series of VAF at a final XNA concentration of 500 nM. Our data showed that at as low as 0.039% VAF, both the mutation and WT peaks were detected with Sanger sequencing (Fig. 3b).

**Figure 3.**
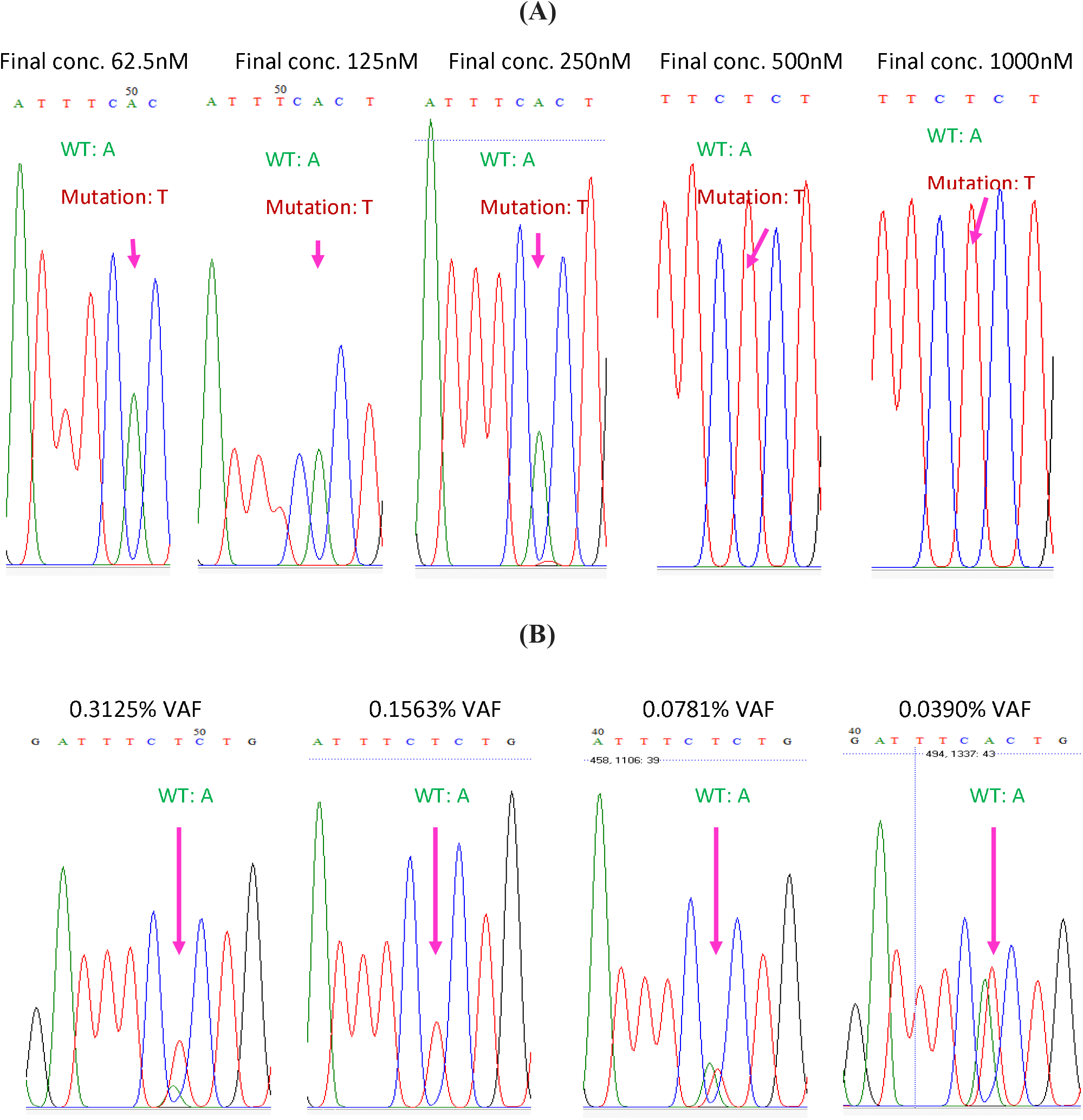
(A). The clamping effect of different BRAF XNA concentration on mutation detection at 0.625% VAF. The reverse primer was used for Sanger sequencing and the reverse/complement strand was read. The c1799 BRAF V600E mutation, the T to A mutation, was read as A to T mutation in sequencing results. A was read as the wildtype and T as the mutant. (B) The limit of detection of BRAF V600E detection in the presence of 500 nM XNA using templates with different VAF. The reverse primer was used for Sanger sequencing and the reverse/complement strand was read. The c1799 BRAF V600E mutation, the T to A mutation, was read as A to T mutation in sequencing results. A was read as the wildtype and T as the mutant.

### The BRAF V600 XNA also enriches the BRAF V600K and V600R mutations

It is important to know if the XNA used to detect the BRAF V600E mutation can also enrich other clinically important BRAF V600 variants, such as V600K and V600R. We used 10 ng 1.25% VAF of BRAF V600K and V600R mutations as the templates and performed the PCR in the presence of BRAF V600 XNA at a final concentration of 500 nM. Both BRAF V600K and V600R mutations were detected (see Figure 4), indicating that the BRAF V600 XNA can also enrich V600K and V600R mutations for detection by Sanger sequencing at least at 1.25% VAF.

**Figure 4.**
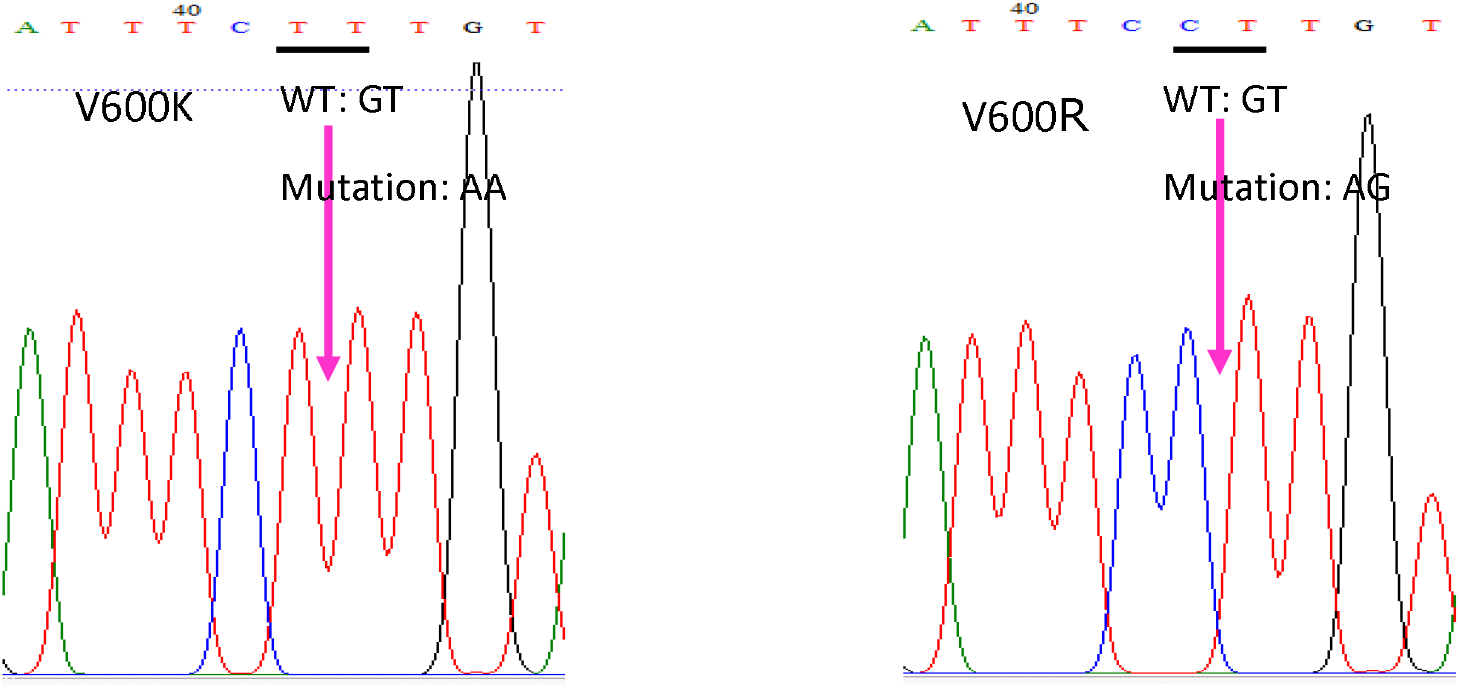
The BRAF V600K and V600R mutations were also detected in the presence of 500 nM XNA using templates with 1.25% VAF. The reverse primer was used for Sanger sequencing and the reverse/complement strand was read. The c1799 BRAF V600K mutation, the GT to AA mutation, was read as AC to TT mutation in the sequencing result (left). The c1799 BRAF V600R mutation, the GT to AG mutation, was read as AC to CT mutation in the sequencing result (right).

### BRAF V600E mutation enrichment and detection in clinical samples

To verify that the BRAF V600E mutation can be enriched by BARF V600 XNA in clinical samples, we tested three BRAF V600E-positive (#82, 83, and 86) and one BRAF V600E-negative (#87) clinical samples. The genomic DNA was isolated from FFPE samples and 10 ng DNA was used as the template for the BRAF V600E mutation enrichment PCR in the presence or absence of BRAF V600 XNA and subsequent Sanger sequencing. The Sanger sequencing result shows that BRAF V600E was detected in all these positive samples where BRAF V600 XNA was added in the PCR, but not in the negative sample (Fig. 5).

**Figure 5.**
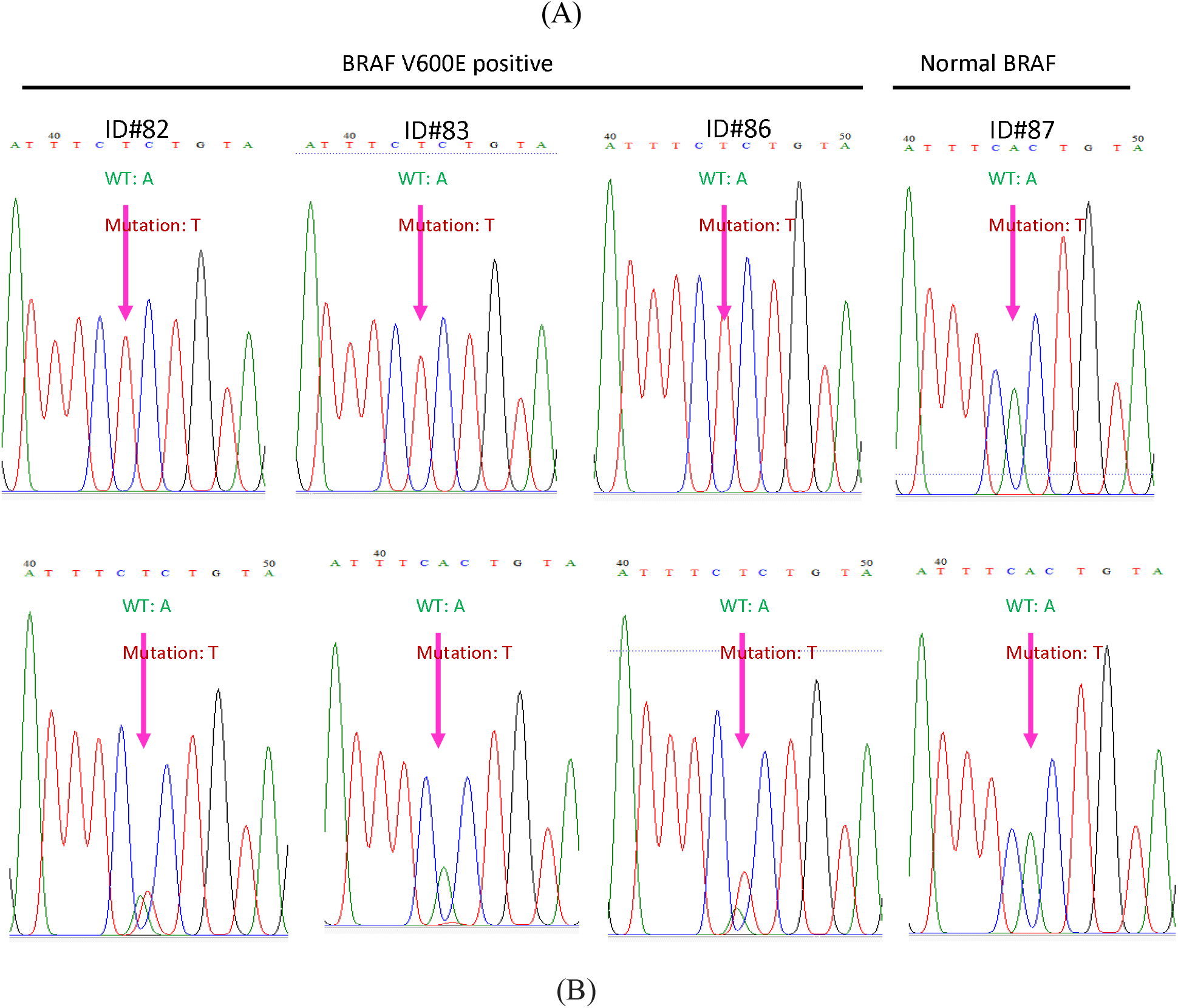
The BRAF V600E mutation was detected in clinical samples. (A) PCR with BRAF XNA, (B) PCR without BRAF XNA. The reverse primer was used for Sanger sequencing and the reverse/complement strand was read. The c1799 BRAF V600E mutation, the T to A mutation, was read as A to T mutation in sequencing results. A was read as the wildtype and T as the mutant.

We found that for clinical samples # 82 and #86, both the mutation and wildtype peaks were detected when no BRAF V600 XNA was added, suggesting the VAF of these samples are more than the Sanger sequencing detection limit, which is 20%. However, only the mutation peak was detected after BRAF V600 XNA was added in these samples, suggesting that the wildtype sequence amplification in these samples had been totally blocked by the XNA. For sample #83, the mutation peak was not detected without the presence of BRAF V600 XNA, suggesting that the VAF is probably less than 20%; when the BRAF V600 XNA was added, only the mutant peak was seen, again suggesting that the amplification of the wildtype sequence had been inhibited. For the negative sample, only the wildtype peak was detected regardless of the presence of the BRAF V600 XNA.

### Validation Protocol for BRAF V600 Mutation Enrichment-Coupled Sanger Sequencing

We have shown that target enrichment has improved the Sanger sequencing sensitivity. To create a validation protocol for BRAF V600 mutation enrichment-coupled Sanger Sequencing that any clinical labs can use, the BRAF V600 mutation was enriched by the BRAF V600 XNA and Sanger sequencing was conducted for downstream validation. We used Sanger sequencing kits manufactured at AdvancedSeq as described in the method section to create this protocol. The Sanger sequencing was conducted using both the forward and reverse primers.

We first sequenced the two positive controls and the negative control. The two positive controls contained the BRAF V600E mutant DNA at 1.25% and 5% VAF, respectively, while the negative control contained only the wildtype DNA (Figure 6). Our results showed that the BRAF V600E mutations could be detected in both positive controls in the presence of XNA as a single mutation peak, but only the wildtype peak was identified in the absence of XNA. These results indicate that in the presence of BRAF V600E XNA, the V600E mutation can be detected at as low as 1.25% VAF. The negative control, however, does not detect the mutation regardless of the presence of the XNA.

**Figure 6.**
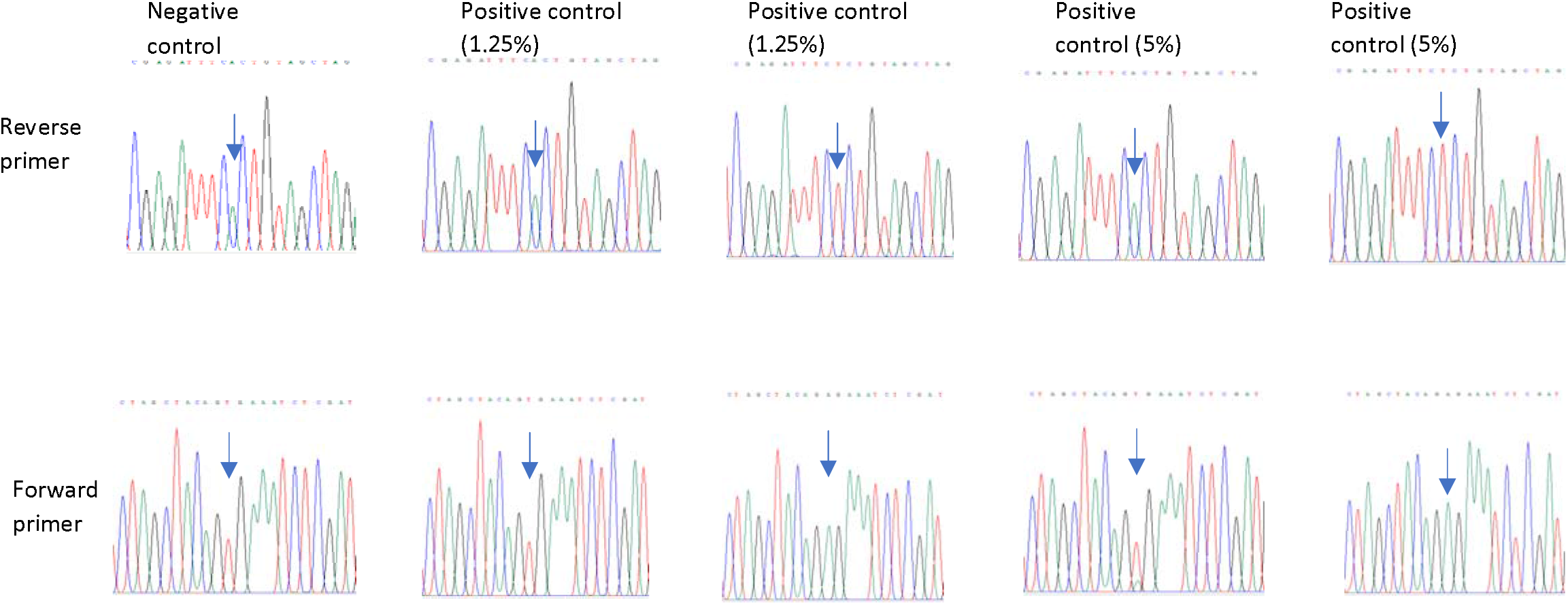
Sequencing of the positive controls (BRAF V600E mutant with 1.25% and 5% VAF) and negative controls (0% mutant) for the BRAF V600E mutations. The negative control shows the wildtype sequence while the BRAF V600E mutation in positive controls are identified after enrichment in the presence of BRAF V600 XNA followed by Sanger sequencing using both forward and reverse primers. The arrows point to the base where the mutation can occur. For forward primer sequencing, the wildtype reads T, and the mutant reads A; for reverse primer sequencing, the wildtype reads A, and the mutant reads T.

We also tested the three clinical samples (BRAF-positive #82, #86, and BRAF-negative #87) and found all these V600E mutations were detected in the BRAF positive samples but not in the normal BRAF sample (data not shown), with similar results shown in the feasibility test above.

## Discussion

Different testing methods have been developed for BRAF V600E detection, including the FDA-approved tests for BRAF V600E mutations as companion diagnostic tests for the use of inhibitors in different types of cancers (25). The early BRAF V600E mutation detection test approved by the FDA uses highly sensitive qPCR method. The Cobas 4800 BRAF V600 Mutation Test, the earliest CDx test approved by the FDA for BRAF V600 mutation in melanoma using real-time PCR method, accurately detects BRAF V600E, but fails to detect BRAF V600K consistently (26, 27, 28). Just like BRAF V600E, V600K is also an important mutation responsive to vemurafenib. The THXID BRAF kit uses ARMS-real time PCR to detect BRAF V600E mutation for dabrafenib and trametinib treatment and BRAF V600K mutation for trametinib treatment. Qiagen’s Therascreen BRAF V600E RGQ OCR kit has better sensitivity (1.87% VAF) compared to the Cobas 4800 BRAF V600 Mutation Test (5% VAF) and the THXID BRAF Kit (5% VAF), and it can detect different V600 mutations.

Droplet digital PCR (ddPCR) technology, whose high sensitivity is claimed to reach 0.001% VAF, has also been used to detect BRAF V600E mutation (21, 29). Albeit its high sensitivity, the ddPCR method also suffers from compromised specificity (with high false-positive results). Although the assay can be optimized to overcome the specificity issue (30), ddPCR for BRAF V600E mutation detection can miss V600K mutation (31). In addition, like other qPCR assays, ddPCR assays can only detect mutations but could not delineate the nature of the mutation as sequencing technologies can.

NGS is a powerful tool when multiple mutations in different genes need to be detected in a single assay. Multiple gene mutations can be involved in cancers such as non-small cell lung cancer (NSCLC), and NGS uses a targeted panel to detect mutations in all those genes, helping physicians make optimal decision at the first line of treatment. The two FDA-approved NGS tests that cover the BRAF V600 mutations include Foundation Medicine’s FoundationOne CDx (covering 324 genes) and Thermo Fisher Scientific’s Oncomine Dx Target Test (covering 23 genes). These panels complement the traditional molecular diagnostics techniques and enable multiple gene mutation detection for cancer diagnostics. But for BRAF V600 mutation detection alone, these NGS-based assays are more time-consuming and less cost-effective compared to Sanger sequencing.

Traditional Sanger sequencing provides accurate sequencing results and is the gold standard for mutation detection, but its major limitation lies in its low sensitivity. The method requires samples with mutant/normal cell population ratio, namely VAF, to be over 15 to 20% (32). Such a low sensitivity hampers Sanger sequencing from being the first line companion diagnostic method for target therapies of important cancer mutations, including the BRAF V600E mutation test.

Efforts to improve the sensitivity of Sanger sequencing have been made either by improving the analytical software (33) or by adding a wildtype blocking agent such as LNA to the PCR reactions that are used for preparing Sanger sequencing templates (34, 31, 35). The software solution improves the sensitivity to 5% from 20%. The wildtype blocking agents improve sensitivity to 0.2 to 0.7%. The blocking reagent can also help improve NGS sensitivity by reducing the sequencing depth (36, 37).

Here we report that we can improve Sanger sequencing sensitivity to 0.04% VAF using our proprietary XNA technology. The XNA molecular clamps method adds XNA molecular clamps to regular PCR reactions to block the wildtype (normal) DNA sequence amplification and to selectively amplify the mutant sequence (38). The use of XNA significantly increases assay sensitivity and helps detect low-frequency mutations at 0.1 to 0.01% VAF (19, 20), which is much lower than the standard qPCR sensitivity (about 1% VAF).

To directly detect the BRAF V600E mutation, we combined the XNA technology with Sanger sequencing because of the ease with which it can be applied in any genomic clinical lab where capillary sequencing facility is common. We have demonstrated that the test can be validated in individual CLIA labs for BRAF V600E testing as a potential companion diagnostic test using commercial Sanger sequencing kits. The validation protocol described in this paper can be easily adapted in other labs. For a better sensitivity test, the two positive controls set at 1.25% or 5% can also be adjusted at a lower VAF if needed for validation. The XNA-based Sanger sequencing method does not add any additional steps to standard Sanger sequencing. The only difference is the enrichment of the mutation with XNA in the PCR step when preparing the template. The mutation-enrichment coupled Sanger sequencing method can be carried out in routine labs with a Sanger sequencing facility and can be the preferred testing technique in labs that are budget-constrained for other expensive instruments.

The XNA-based method is also comparable if not better than other blocking reagents with regards to its blocking capability. For example, our XNA has better blocking capability for KRAS compared to the LNA-based blocking reagent (34), blocking KRAS G12 mutation at 0.02% VAF (data not published) while the LNA-based KRAS blocking reagent does not block the wildtype amplification until 10% VAF of the mutation sequence is present. In lower concentrations the wildtype sequence amplification is not blocked but they were able to amplify and detect the mutant sequence down to 0.7%. For our XNA-based blocking agent, the wildtype sequence amplification is either totally blocked or partially blocked between 0.3 to 0.04% VAF of the BRAF mutation, but the mutant sequence is strongly amplified as the only peak or a peak with similar height of the wildtype peak.

In summary, in this paper we present a new method for combining BRAF V600E mutation enrichment and validation using Sanger sequencing for the detection of BRAF V600 mutations with increased sensitivity and without adding additional steps or instruments. This method can significantly improve the sensitivity of Sanger sequencing to 0.04% VAF. Any regular clinical lab should be able to easily validate this method based on our protocol presented here for sequencing the BRAF V600E mutation. Furthermore, this highly sensitive target-enrichment coupled Sanger sequencing technique should work for the detection of other single low-frequency mutations.

